# Phage lysates are reservoirs of biologically active quorum sensing signals

**DOI:** 10.1101/2025.04.11.648453

**Authors:** Amalie Høgh Eichler, Jesper Juel Maruritzen, Kira Céline Koonce, Gagandeep Kaur, Nina Molin Høyland-Kroghsbo

**Author notes:** Address correspondence to Nina Molin Høyland-Kroghsbo.

## Abstract

Bacteriophage lysates are widely used to study phage-bacterial interactions and are typically applied after filtration without further purification. As a result, host-derived metabolites produced during phage propagation are transferred alongside phage particles, yet the biological consequences of this carryover remain largely unexplored. Here, we show that the *Pseudomonas aeruginosa* quorum sensing signal PQS persists in a bioactive state in phage lysates for at least one year. In contrast, the quorum sensing molecules 3OC12-HSL and C4-HSL are rapidly degraded. We demonstrate that PQS stability is caused by its association with outer membrane vesicles, which protect the signal from degradation. PQS from phage lysates induces production of the virulence factor pyocyanin and increases mortality in a *Galleria mellonella* model of *P. aeruginosa* infection. Importantly, PQS carryover does not explain previous reports of phage-mediated activation of the PQS system, as this response also occurs in PQS-free phage lysates, establishing phage lysate-mediated signal transfer and phage-induced signaling as distinct mechanisms. Because hydrophobic signaling molecules are associated with outer membrane vesicles in diverse Gram-negative bacteria, including *Vibrio harveyi* (CAI-1) and *Paracoccus* (C16-HSL), our findings suggest that phage lysates may serve as reservoirs and overlooked vectors of host-derived communication signals in a broad range of phage-host systems. Our results reveal a previously overlooked layer of phage-host interactions in which biologically active bacterial signals persist beyond phage lysate preparation and influence recipient-cell behavior independently of phage infection.

**Significance:** Bacteria coordinate collective behaviors including virulence through chemical communication. We show that one such signal, PQS, remains biologically active in phage lysates because it is protected within outer membrane vesicles. Specifically, lysate-associated PQS alters bacterial physiology and inadvertently increases virulence in an infection model. We identify phage lysates as previously unrecognized reservoirs of host-derived communication signals, revealing a mechanism by which bacterial signaling activity can persist beyond phage lysate production and potentially confound phage-host interaction studies across diverse bacterial species. This phenomenon may also have practical consequences for phage therapy development, as lysate-associated signaling molecules could mask phage efficacy by transiently enhancing bacterial virulence, potentially causing promising therapeutic candidates to be discarded during early-stage evaluation.

## Introduction

Bacteriophage (phage) viruses infect bacteria and shape microbial communities and behavior. Over the past decade, intricate interactions between bacterial cell-to-cell signaling, known as quorum sensing (QS), and outcomes of phage-bacterial encounters have been uncovered. QS is the process that allows bacteria to communicate and coordinate collective behaviors through the production and detection of signaling molecules, also known as autoinducers, in response to changes in population density (Papenfort & Bassler, 2016). Since QS molecules accumulate in step with increasing cell density, and given that phages spread more rapidly when susceptible host cells are densely packed, bacteria can use QS levels at high cell density as an indicator of high risk of phage infection and regulate their anti-phage defenses, including receptor expression, CRISPR-Cas activity, and defense strategy selection (Høyland-Kroghsbo et al., 2013, 2017; Patterson et al., 2016; Tan et al., 2015). Conversely, some phages can directly sense or manipulate host QS systems to optimize their fitness (Erez et al., 2017; Schwartzkopf et al., 2023; Shah et al., 2021; Silpe & Bassler, 2018). Together, these studies highlight the central role of QS and QS inhibition in shaping phage-bacterial interactions.

In *Pseudomonas aeruginosa*, QS systems are generally considered to be orchestrated hierarchically. First, the synthase LasI produces the 3-oxo-C12-homoserine lactone (3OC12-HSL), activating its cognate receptor LasR, which turns on transcription of virulence genes and additionally activates the RhlR/I system (Latifi et al., 1995; Pesci et al., 1997; Seed et al., 1995). The synthase RhlI produces C4-homoserine lactone (C4-HSL), which activates the RhlR receptor that also regulates virulence genes (Pearson et al., 1995). 3OC12-HSL and C4-HSL belong to the acyl-homoserine lactone (AHL) type of QS molecules, which consist of a homoserine lactone moiety and a variable acyl side chain, and are commonly produced by Gram-negative bacteria (Papenfort & Bassler, 2016). The Las and Rhl systems together, in opposite manners, control the PQS QS system (Gallagher et al., 2002; Wade et al., 2005; Xiao, He, et al., 2006). The enzymes PqsABCD synthesize 2-heptyl-4-quinolone (HHQ) (Déziel et al., 2004; Dulcey et al., 2013), which, in a final PqsH-controlled step, is converted into 2-heptyl-3-hydroxy-4-quinolone (PQS) (Schertzer et al., 2010).

Interestingly, phage infection activates the PQS signal response in *P. aeruginosa* (Blasdel et al., 2018; Bru et al., 2019; Høyland-Kroghsbo & Bassler, 2022). Specifically, in the absence of *lasI*, phage infection overrides the hierarchical QS network and activates the PQS response, inducing production of the PQS-controlled virulence factor pyocyanin (Høyland-Kroghsbo & Bassler, 2022). Furthermore, phage infection causes *P. aeruginosa* colonies to emit PQS. Uninfected *P. aeruginosa* swarms sense the PQS stress signal and change swarming direction, thereby avoiding the infection site (Bru et al., 2019).

Another PQS-mediated defense strategy is the increased production of membrane vesicles. Due to its hydrophobicity, PQS intercalates between the lipopolysaccharides of the outer membrane, creating curvature of the membrane and formation of outer membrane vesicles (Mashburn-Warren et al., 2008). These vesicles simultaneously serve as delivery vehicles for PQS within the *P. aeruginosa* population (Mashburn & Whiteley, 2005) and as phage decoys, depleting the phage population (Augustyniak et al., 2022; Manning & Kuehn, 2011; Reyes-Robles et al., 2018). Thus, outer membrane vesicles play a complex role in phage-bacterial dynamics.

Filter sterilized phage lysates contain not only phages but also host-derived material released during lysis, including outer membrane vesicles and signaling molecules. As phage particles and outer membrane vesicles are similar in size (approximately 20 to 200 nm), both are commonly retained during standard lysate preparation (Paczesny & Bielec, 2020). Consequently, host-derived factors are transferred alongside phages, yet their bioactivity and contribution to fundamental phage-host interaction studies remain poorly understood.

Here, we show that the *P. aeruginosa* QS molecule PQS persists in phage lysates through stabilization within outer membrane vesicles. These findings reveal a previously overlooked mechanism by which host-derived signaling molecules can influence bacterial phenotypes and potentially confound the outcomes of phage-host interaction studies.

## Results

### The QS molecule PQS is abundant in phage lysates

We initially set out to investigate the timing and the underlying mechanism of the phage-induced PQS induction in *P. aeruginosa* UCBPP-PA14 (PA14) (Bru et al., 2019; Høyland-Kroghsbo & Bassler, 2022), using the virulent phage DMS3^vir (^Bru et al., 2019). To avoid confounding effects of the DMS3-encoded LasR inhibitor Aqs1, we generated a DMS3^vir^ *^a^qs1* mutant as in (Shah et al., 2021)(Shah et al., 2021), which we denote DMS3^vir-*aqs1*.^ The mutant produces an Aqs1 protein with an amino acid substitution that abolishes Aqs1 binding to LasR (Shah et al., 2021). Surprisingly, we detected a PQS signal in this phage lysate control before adding cells (Supplementary Fig. 1), indicating that PQS produced during lysate production on the propagation PA14 was present in the lysate.

This observation led us to investigate PQS levels in phage lysates. Therefore, we constructed an *Escherichia coli* PQS reporter strain expressing the PQS receptor PqsR from pBAD and carrying the PqsR-activated *pqsA* promoter in front of *luxCDABE* in pCS26 as described in (Paczkowski et al., 2017).

We prepared new phage lysates on PA14 Δ*pf5* (denoted QS+ lysates) or on a triple QS-synthase mutant PA14 Δ*pf5* Δ*lasI* Δ*rhlI* Δ*pqsA* (denoted QS-lysates). We selected the Δ*pf5* background to eliminate potential interference from the prophage Pf5 residing in the PA14 genome, since the related *P. aeruginosa* PAO1 prophage Pf4 interferes with PQS synthesis (Schwartzkopf et al., 2023)(Schwartzkopf et al., 2023). Using our PQS reporter, we found that PA14 ON culture supernatants contained an average of 9.60 µM PQS (Fig. 1A), consistent with previously reported values of 5-11 µM (Bala et al., 2013; Diggle et al., 2003)(Bala et al., 2013; Diggle et al., 2003). We were unable to detect PQS in the QS-phage lysates (<0.002 µM), as expected, whereas QS+ lysates contained an average of 32.0 µM PQS (Fig. 1A). These data demonstrate that PQS produced by the propagation host is transferred to and remains abundant in filter-sterilized phage lysates.

**Fig. 1.**
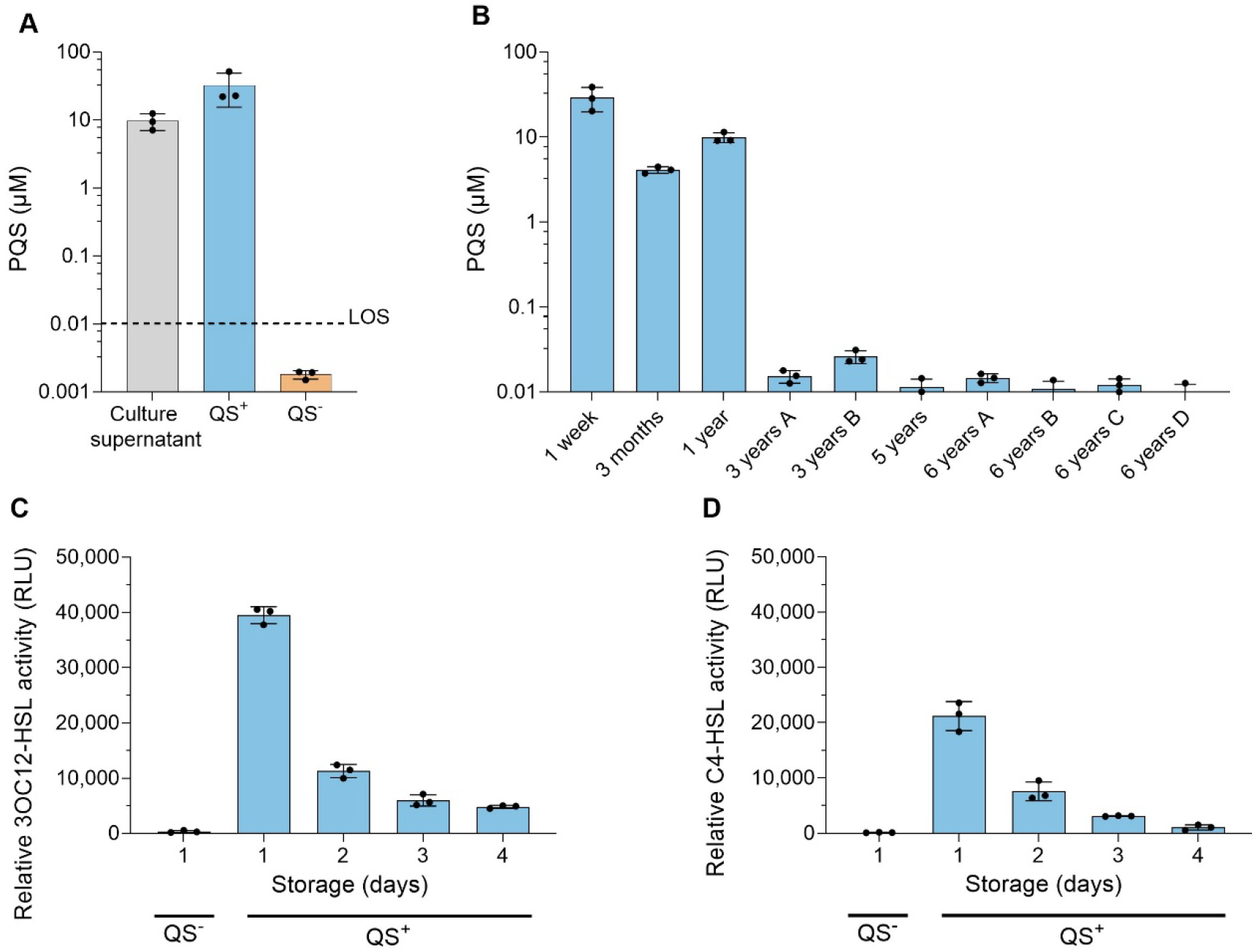
QS signals in phage lysates. A. The concentration of PQS in ON cultures of PA14 and in DMS3^vir-*aqs1* p^hage lysates generated on PA14 Δ*pf5* (QS+ lysate) or PA14 Δ*pf5* Δ*lasI* Δ*rhlI* Δ*pqsA* (QS-lysate), quantified using an *E. coli* PQS reporter. LOS = limit of sensitivity. Error bars represent the standard deviation of three biological replicates. B. PQS concentrations in phage lysates stored between one week and up to six years at 4 °C, measured as in A. The lysates, representing five different phages, were produced either as plate lysates (1 week, 3 months, 1 year, 5 years, 6 years A-D) or by lysogen induction (3 years A-B). Error bars represent the standard deviation of three technical replicates. Y-axis begins at the limit of sensitivity (0.01 µM). B. 3OC12-HSL and C. C4-HSL signals were quantified immediately in freshly prepared QS- and QS+ lysates, and every 24 h for a period of four days for the latter, using *E. coli* 3OC12-HSL or *E. coli* C4-HSL reporters, respectively. Lysates were stored at 4 °C between measurements. RLU indicates the luminescence signal normalized to the OD_600 o_f the 3OC12-HSL and the C4-HSL reporters, respectively. Error bars represent the standard deviation of three biological replicates. B-D. The specifics of the phage lysates are detailed in Supplementary Table 2.

### PQS is stable in phage lysates but AHLs are rapidly degraded

To test PQS stability, we quantified PQS in phage lysates stored at 4 °C for up to 6 years. PQS remained abundant after 1 week (29.2 µM) and persisted at high concentrations after 3 months (4.11 µM) and 1 year (9.93 µM) (Fig. 1B). Even lysates stored for 3 to 6 years retained detectable levels of PQS (10.0-26.2 nM). These findings show that bioactive PQS persists in phage lysates for at least 1 year and can remain detectable for several years.

To assess the stability of the two other major *P. aeruginosa* QS molecules, 3OC12-HSL and C4-HSL, we used previously described *E. coli* reporter strains (Paczkowski et al., 2017). Both 3OC12-HSL and C4-HSL were detected at low levels in the freshly prepared QS+ lysates but were absent from QS-lysates, as expected (Fig. 1C-D). Standard curves generated with synthetic 3OC12-HSL and C4-HSL showed that signals from lysates were below the linear quantification range of the assays (<1 nM and <1 µM, respectively). Therefore, Fig. 1C-D presents the reporter output as relative light units (RLU) rather than concentrations. In contrast to PQS, both AHLs were highly unstable and became nearly undetectable after 3-4 days of storage, consistent with previous findings (Byers et al., 2002; Glansdorp et al., 2004; Yates et al., 2002). These findings indicate that 3OC12-HSL and C4-HSL are rapidly degraded in phage lysates and are therefore unlikely to substantially influence downstream applications unless lysates are used immediately after preparation.

### PQS in phage lysates is bound within membrane vesicles

The hydrophobic QS molecule PQS accumulates in the outer bacterial membrane (Florez et al., 2017), where it causes membrane bending and formation of PQS-containing outer membrane vesicles (Mashburn-Warren et al., 2008). To test how much of the PQS in phage lysates is bound within these vesicles, we filtered phage lysates a second time through 10 kDa filters, which retain outer membrane vesicles, while vesicle-free lysate passes through. Unfiltered QS+ lysates contained 28.3 µM PQS, whereas the vesicle-free QS+ lysates contained 0.016 µM PQS (Fig. 2A). The washed and resuspended outer membrane vesicles contained 23.6 µM PQS. Thus, 83% of the PQS in fresh phage lysates is bound within membrane vesicles, in accordance with previous findings that 86% of the PQS from fresh *P. aeruginosa* cultures is bound within membrane vesicles (Mashburn & Whiteley, 2005).

**Fig. 2.**
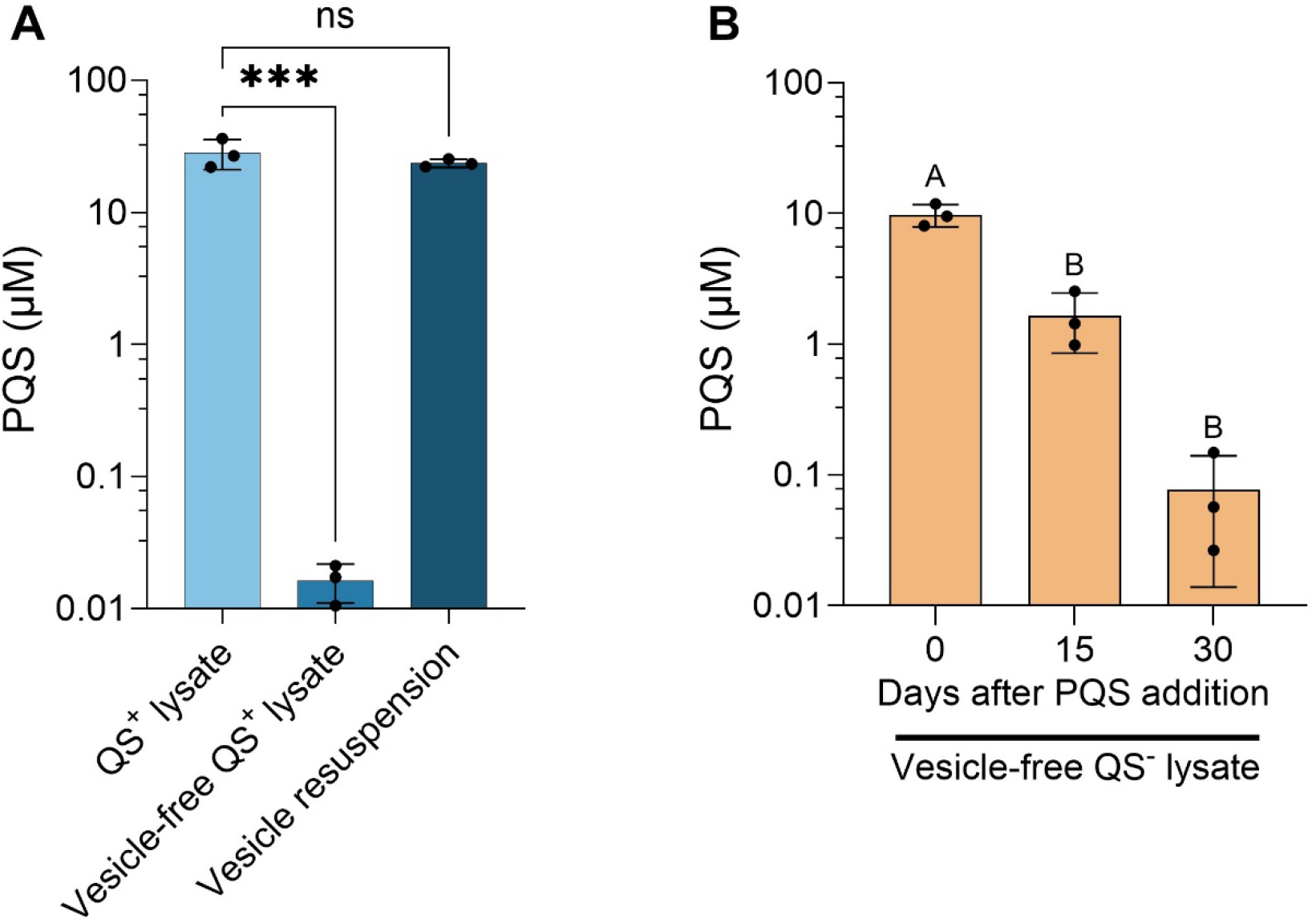
Membrane vesicles provide PQS stability. A. Filter-sterilized DMS3^vir-*aqs1*^ QS+ phage lysate was filtered through a 10 kDa filter, which retains larger molecules, including membrane vesicles and phage particles. The retained particles were washed and resuspended 1:1. The PQS concentration of the QS+ phage lysate, the vesicle-free lysate, and the vesicles was measured as in Fig. 1-2. Error bars represent the standard deviation of three biological replicates. P-values: ns = 0.41, *** = 0.0005 (One-way ANOVA with post-hoc Tukey HSD). B. The stability of synthetic PQS in vesicle-free QS-phage lysates. DMS3^vir-*aqs1*^ QS-lysate was filtered through a 10 kDa filter to remove membrane vesicles. The vesicle-free lysate was supplemented with 10 μM PQS and stored at 4 °C for 0, 15, or 30 days. The PQS concentration of all samples was measured immediately after preparing the last sample as in Fig. 1-2. The signal from a vesicle-free lysate supplemented with DMSO (solvent control) was subtracted from all samples (< 0.02 µM). Error bars indicate the SD of three biological replicates. P = 0.0001. Letters represent statistical groups (one-way ANOVA with post-hoc Tukey HSD). A-B. Y-axes begin at the limit of sensitivity (0.01 µM).

To determine whether PQS stability was caused by its association with the outer membrane vesicles, vesicle-free QS-lysates were generated by 10 kDa filtration and supplemented with 10 μM synthetic PQS. Following storage at 4 °C, PQS concentrations decreased from 9.77 μM at day 0 to 1.65 μM after 15 days and 0.08 μM after 30 days (Fig. 2B). Thus, in the absence of outer membrane vesicles, bioactive PQS declined by more than 100-fold within one month, demonstrating that membrane vesicles are largely responsible for the remarkable stability of PQS in phage lysates.

### PQS transfer via phage lysates induces PQS-regulated virulence factor production

Since phage lysates stored for up to one year contain physiologically relevant concentrations of PQS, we tested the ability of phage lysates to induce production of the PQS-controlled virulence factor pyocyanin. To isolate the effect of PQS transfer from those of phage infection-mediated PQS activation (Blasdel et al., 2018; Bru et al., 2019; Høyland-Kroghsbo & Bassler, 2022), we used a *pilA*::FRT mutant lacking type IV pili. This mutant, hereafter referred to as the *pilA* mutant, is resistant to infection by DMS3^vir-*aqs1*^, which requires type IV pili for infection (Budzik et al., 2004). We also constructed a *pilA* Δ*pqsA* double mutant that is both phage resistant and unable to synthesize PQS, rendering it deficient in pyocyanin production (Xiao, Déziel, et al., 2006).

To determine whether lysate-derived PQS can induce *P. aeruginosa* virulence factor production, we grew the DMS3^vir-*aqs1*-^resistant *pilA* mutant in the absence of phage lysate, as a control to measure baseline pyocyanin production, and the *pilA* Δ*pqsA* double mutant in the presence of 1-10% QS+ or QS-lysate, concentrations commonly used to achieve the desired multiplicity of infection (MOI). As expected, the PQS-proficient *pilA* strain produced pyocyanin, whereas the PQS-deficient *pilA* Δ*pqsA* strain did not (Fig. 3A). Addition of QS+ phage lysate partially restored pyocyanin production in the *pilA* Δ*pqsA* strain in a dose dependent-manner with 10% lysate restoring pyocyanin levels to 60% of those observed in the *pilA* control strain (Fig. 3A-B). In contrast, QS-lysates did not affect pyocyanin levels of the *pilA* Δ*pqsA* strain (Fig. 3A-B). These data demonstrate that PQS present in a standard phage lysates is sufficient to activate PQS-controlled virulence factor production in recipient *P. aeruginosa* cells.

**Fig. 3.**
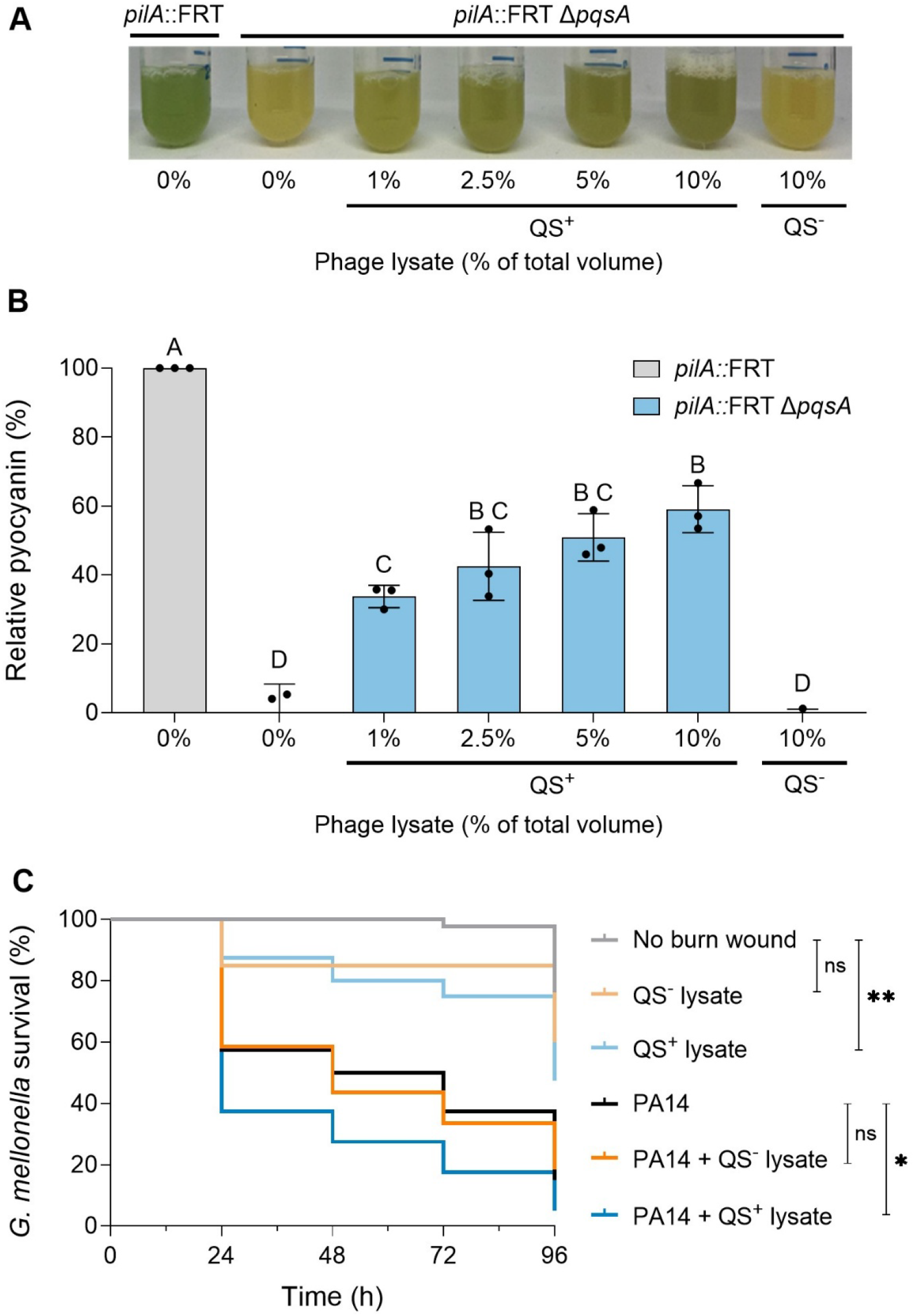
PQS in phage lysates induces *P. aeruginosa* virulence. A+B. The effect of phage lysates on pyocyanin virulence factor production. The DMS3^vir-*aqs1*-^ resistant *pilA::*FRT mutant and *pilA::*FRT Δ*pqsA* mutant were cultured with either control (LB) or up to 10% of 5*10^10 P^FU/mL QS+ or QS-phage lysate, respectively. A. Image of representative cultures. Pyocyanin is visible as a green pigment. B. Relative pyocyanin production of the cultures. Pyocyanin was measured as OD_695 o_f cell-free supernatants, relative to the OD_600 o_f the bacterial cultures. *pilA::*FRT pyocyanin production was set to 100%. Error bars represent the standard deviation of three biological replicates. P-value < 0.0001. Letters represent statistical groups (one-way ANOVA with post-hoc Tukey HSD). C. The effect of phage lysates on *G. mellonella* larvae survival. Larvae were either left untreated or inflicted with a burn wound and treated with QS- or QS+ phage lysate alone, or infected with PA14 WT, incubated for one h, and treated with either QS- or QS+ phage lysate. Larvae survival was monitored over 96 h. Data represent mean survival from three independent experiments (n = 30 larvae per condition). ns = non-significant. No burn wound vs QS+ phage treatment (**, p = 0.0027), PA14 vs PA14 + QS+ phage lysate. (*, p = 0.0291) Statistical analysis was performed using a log-rank test based on end-point mortality. However, it should be noted that the *pqsABCDE* activity between 4 and 8 h is significantly enhanced in response to phage infection from the QS+ lysate, compared to both the uninfected control (***, p = 0.0002), and to QS-lysate infection (****, p <0.0001), underlining that lysate-derived PQS transiently amplifies phage-induced PQS response (Fig 4B insert).

To investigate the *in vivo* relevance of lysate-mediated PQS transfer, we used a *Galleria mellonella* burn wound *P. aeruginosa* infection model (Maslova et al., 2020). Larvae were either left untreated or inflicted with a burn wound and immediately treated with QS− or QS+ phage lysate alone or infected with PA14. For the latter group, burn wounds were additionally treated with the respective phage lysates one hour after infection. Survival was monitored over the course of 96 h.

As expected, untreated larvae exhibited high survival throughout the experiment (Fig. 3C). Burn wounds treated with QS− phage lysate alone did not significantly affect survival compared to untreated controls (p = 0.2379), while QS+ lysate treatment alone caused a significant reduction in survival (**, p = 0.0027), indicating that lysate-derived PQS and/or associated components can exert a virulence effect even in the absence of bacterial infection. Even though the DMS3^vir-*aqs1* p^hage effectively kills PA14 *in vitro*, it did not improve survival in this experiment. Infection with PA14 followed by treatment with QS− lysate resulted in moderate mortality, which was not significantly different from PA14 infection alone (p = 0.9008). In contrast, PA14 infection followed by QS+ lysate significantly reduced larval survival compared to PA14 infection alone (*, p = 0.0291), as well as compared to PA14 treated with QS− lysate (*, p = 0.0488). Collectively, these results demonstrate that PQS transferred in QS+ phage lysates can exert virulence in the absence of bacterial infection and significantly enhance virulence during *P. aeruginosa* infection. These results emphasize the importance of considering PQS in phage preparations, as lysate-derived signaling molecules can confound both *in vitro* and *in vivo* studies of phage-bacterial interactions and worsen therapeutic efficacy.

### PQS-free phage lysates induce the PQS stress response

Since phage treatment induces PQS production in both WT PA14 (Bru et al., 2019) and in a *lasI* mutant (Høyland-Kroghsbo & Bassler, 2022), we asked whether this induction could be explained by PQS present in the phage lysates. To address this, we created a chromosomal *pqsABCDE*::*lux* transcriptional reporter strain (hereafter referred to as the PA14 PqsA reporter) to monitor PQS-circuit activity. We treated the PA14 PqsA reporter with 10% QS+ or QS-phage lysates, corresponding to an MOI of 10, and monitored bacterial growth and relative *pqsABCDE* expression over time.

Fig. 4A shows that both QS+ and QS-phage lysates caused similar levels of bacterial killing, compared to the uninfected control. Despite the absence of PQS in QS-lysates, both QS+ and QS-lysates induced strong activation of *pqsABCDE*, as indicated by the higher relative luminescence in phage-infected samples compared with the uninfected control (Fig. 4B). Notably, the infected cultures show even stronger signals despite their substantially lower OD_600 v_alues, a condition in which PQS-dependent QS signaling is typically not observed. These data show that phage infection itself induces PQS circuit activation and cannot be explained solely by PQS transferred in the lysate. Similarly, infection with both QS+ and OS-phage lysates cause PA14 swarm repulsion (Supplementary Fig. 2), consistent with previous findings (Bru et al., 2019).

**Fig. 4.**
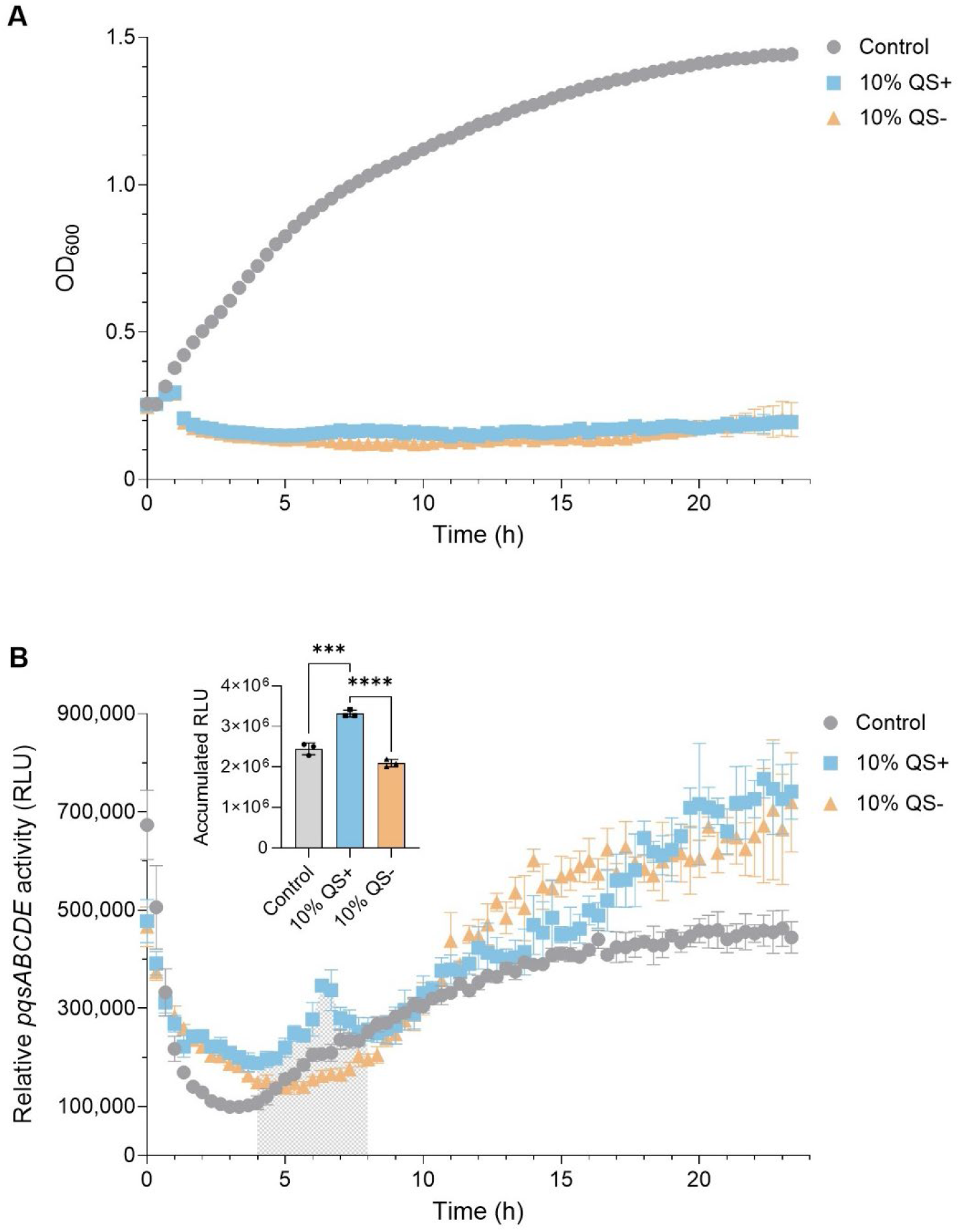
Phage infection from PQS-free lysates induces PQS activation. The effect of QS+ or QS-phage lysates on PQS QS circuit activity measured using a PA14 *pqsABCDE* reporter. The reporter strain was treated with 10% LB or QS+ or QS-phage lysate at an OD_600 o_f 0.5, equivalent to an MOI of 10 for phage infection. A. Bacterial growth measured as OD_600 o_ver time. B. The relative *pqsABCDE* activity was assessed as RLU (luminescence/OD_600)_. The insert shows the accumulated RLU of each treatment between 4-8 (the grey area). Error bars represent the standard deviation of three biological replicates. P-values: *** = 0.0002, **** = <0.0001 (One-way ANOVA with post-hoc Tukey HSD). A-C.

Together, our data show that vesicle-bound PQS transferred in phage lysates can induce virulence factor production in the recipient *P. aeruginosa* cultures *in vitro* and enhances virulence *in vivo,* potentially a confounding factor in studies of phage-bacterial interactions and phage efficacy. Importantly, this effect is distinct from phage-induced activation of the PQS system. Infection with PQS-free phage lysates also induced *pqsABCDE* expression in PA14, confirming that phage infection itself activates PQS, in accordance with previous observations (Bru et al., 2019; Høyland-Kroghsbo & Bassler, 2022). Collectively, these findings establish phage lysate-mediated PQS transfer and phage-induced PQS activation as distinct yet overlapping mechanisms that should be considered separately when studying *P. aeruginosa* responses to phage infection.

## Discussion

Here, we demonstrate that the *P. aeruginosa* QS molecule PQS persists at biologically relevant concentrations in phage lysates for up to 1 year. PQS present in standard lysates is sufficient to induce PQS-dependent gene expression and enhance virulence *in vivo*. Our findings highlight that phage studies using unpurified *P. aeruginosa* phage lysates may measure bacterial responses to transferred PQS in addition to phage-mediated effects. These confounding effects are likely more pronounced when larger volumes of lysate are added, such as in studies using low-titer-lysates and/or high MOIs. These effects can be mitigated by propagating phages on a QS-deficient host or by purifying the phages prior to use.

In contrast to PQS, the AHL QS molecules 3OC12-HSL and C4-HSL were highly unstable in phage lysates during storage (Fig. 1C-D), consistent with prior reports of fast non-enzymatic turnover of AHLs in a temperature-, and acyl chain length-dependent manner, due to pH-driven hydrolysis of the lactone moiety (Byers et al., 2002; Yates et al., 2002). Neither 3OC12-HSL nor C4-HSL binds within outer membrane vesicles. We show that the exceptional stability of PQS is largely explained by its association with outer membrane vesicles, as removal of vesicles reduced PQS bioactivity by 100-fold within one month (Fig. 2B), suggesting that outer membrane vesicles provide a stabilizing scaffold for hydrophobic QS molecules.

Importantly, PQS is not the only outer membrane vesicle-bound QS molecule. The hydrophobic QS molecules including CAI-1 from *Vibrio harveyi* and long-chain AHLs from *Paracoccus* are also associated with membrane vesicles (Brameyer et al., 2018; Morinaga et al., 2020). These observations raise the possibility that lysate-mediated transfer of stable QS signals may extend beyond *P. aeruginosa* and represent a broader feature of host-phage systems, which may inadvertently affect the results of phage-bacterial interactions. Cell-free phage synthesis has been proposed to minimize potentially bioactive molecules such as endotoxins, which may adversely affect patient outcomes (Turner et al., 2024), while also preventing carryover of host-derived QS signals into phage lysates.

Indeed, we found that PQS present in a standard phage lysate is sufficient to activate PQS-controlled virulence factor production in recipient *P. aeruginosa* cultures (Fig. 3A+B). Similarly, prior research shows that addition of PQS-containing outer membrane vesicles increases the production of multiple virulence- and biofilm-related factors, including pyocyanin, elastase, siderophores, alginate, and rhamnolipids (Bala et al., 2015). Thus, PQS and other vesicle-associated factors likely contribute to the increased mortality observed in QS+ lysate-treated *P. aeruginosa-*infected *G. mellonella* (Fig. 3C). PQS itself is a virulence factor that promotes the generation of reactive oxygen species, causing oxidative stress and tissue damage (Abdalla et al., 2017), while PQS-induced outer membrane vesicles contain a variety of additional virulence-associated factors including PQS (Mashburn & Whiteley, 2005; Rueter & Bielaszewska, 2020). The presence of virulence-factor-containing outer membrane vesicles may be the reason that addition of QS+ lysates enhances mortality even in the absence of a bacterial infection (Fig. 3C). Together, these findings emphasize that both PQS and associated outer membrane vesicles may confound results of phage lysates *in vivo* by prematurely turning on virulence factor production and disturbing the QS machinery of the cell.

While these findings indicate that lysate-derived PQS contributes substantially to increased virulence, they do not imply that PQS carryover is the only mechanism by which phage treatment can influence infection outcomes. Notably, QS-phage treatment of PA14-infected *G. mellonella* did not prevent the larvae from succumbing to the infection (Fig. 3C). While the DMS3^vir-*aqs1* p^hage effectively kills PA14 *in vitro*, this does not translate into enhanced survival *in vivo* in our assay. This may be because phage infection activates the PQS circuit, which in turn enhances virulence factor production (Blasdel et al., 2018; Bru et al., 2019; Høyland-Kroghsbo & Bassler, 2022). Indeed, when we engineered DMS3^vir-*aqs1* t^o produce the PQS-degrading enzyme AqdC, it enhanced survival of PA14-infected *G. mellonella* (Mauritzen et al., 2026).

Although PQS was likely present in the phage lysates used in previous studies on phage-mediated PQS activation (Blasdel et al., 2018; Bru et al., 2019; Høyland-Kroghsbo & Bassler, 2022), our results show that lysate-derived PQS is not the sole driver of virulence modulation. Phages propagated on both QS+ and QS− hosts induced *pqsABCDE* expression upon infection (Fig. 4B), indicating that activation of the PQS circuit is an intrinsic *P. aeruginosa* response to phage infection and may contribute to phage-mediated increases in virulence.

In summary, we demonstrate that phage lysates are reservoirs of biologically active outer membrane vesicle-associated QS molecules. In *P. aeruginosa*, lysate-derived PQS persists for extended periods, alters bacterial physiology, and enhances virulence independently of phage infection. Our findings establish lysate-mediated QS molecule transfer and phage-induced QS signaling as distinct mechanisms and highlight the need to consider host-derived metabolites when interpreting phage-host interactions. More broadly, the association of hydrophobic QS molecules with outer membrane vesicles in diverse bacterial species, including *Paracoccus* and *Vibrio harveyi*, suggests that lysate-mediated transfer of bacterial communication signals may represent a widespread and previously overlooked component of phage-host systems. Together, these findings reveal a new dimension of phage-host interactions in which biologically active bacterial signals persist beyond host-cell lysis and continue to influence recipient-cell behavior.

## Methods

### Bacterial strains, plasmids, and phages

Bacterial strains, plasmids, and phages are listed in Table S1 in the supplemental material. Bacteria were grown at 37 °C with shaking in Lysogeny Broth (LB) broth (1% tryptone, 0.5% yeast extract, 1% NaCl) or on LB agar plates solidified with 15 g agar/L. The *E. coli* reporter strains were grown in LB supplemented with 100 µg/mL kanamycin, 100 µg/mL ampicillin, and 0.1% arabinose.

We cloned an *E. coli* PQS reporter strain, with the PQS receptor PqsR produced from a pBAD plasmid and with the *pqsA* promoter (255 bp upstream of *pqsA*) containing a PqsR binding site (Wade et al., 2005) cloned in front of the *luxCDABE* genes in pCS26, as described in (Paczkowski et al., 2017).

The *pqsABCDE*::*luxCDABE* strain was constructed by PCR amplification of 1000 bp upstream and downstream of the desired insertion site. The *luxCDABE* operon was amplified from the pCS26 plasmid. PCR fragments were assembled into the pEXG2 vector (Rietsch et al., 2005) using NEBuilder HiFi DNA Assembly (NEB#E2621S), placing the *lux* operon three base pairs downstream of *pqsE*. This construct was transformed into *E. coli* SM10λpir and subsequently mobilized into PA14 by conjugation. The PA14 *pilA*::FRT Δ*pqsA* mutant was constructed as in (Høyland-Kroghsbo et al., 2017), except that mutants were selected on LB medium without NaCl, containing 15% (wt/vol) sucrose. Candidate mutants were confirmed with PCR and sequencing.

### Phage lysates

50 µL ON culture of the propagation host was mixed with 50 µL phage stock. 5 mL 50 °C molten top LB agar (with 10 mM MgSO_4,_ 6g/L agar) was added, and the suspension was poured onto a room temperature LB plate. The plate was incubated ON at 37 °C. The following day, 5 mL SM buffer (200 mM NaCl_2,_ 10 mM MgSO_4,_ 50 mM Tris-HCl, pH 7.5) was layered on top of the plate and left to incubate for 4h at RT. The SM buffer was centrifuged at 1874 x g for 10 min to remove debris. The supernatant was filter sterilized using a Steriflip® Vacuum Tube Top Filter (Millipore®, SE1M003M00) with a pore size of 0.45 µm and stored at 4 °C in falcon tubes. The titer of the lysate was determined by serial dilution in SM buffer, followed by plaque assays. Two phage lysates from the PQS stability assays were prepared by growing the lysogenic host strain ON and filter sterilizing the supernatants, as outlined in Supplemental Table 2.

### *E. coli* reporter assays

ON cultures of the *E. coli* reporters were diluted 100x in LB containing 100 µg/mL ampicillin, 50 µg/mL kanamycin, and 0.1% arabinose. The mixture was transferred to a Costar® black clear bottom 96-well microtiter plate (Corning®) and samples were added to a final volume of 200 µL, as appropriate. The OD_600 a_nd luminescence were measured every 20 min for 24 h at 30 °C, with shaking before each measurement using a Synergy H1 plate reader (BioTek Instruments Inc.) and software Gen5 v.3.05.1. The luminescence signals were divided by the corresponding OD_600 t_o obtain RLUs. The RLU signals from 30 min before and after the maximum value were averaged for each sample to obtain representative single values. Synthetic PQS, 3OC12-HSL, and C4-HSL dissolved in DMSO were included in each plate. The average RLU were log-transformed, and a linear regression model was fitted to the standard curve, and the coefficients were used to calculate the concentrations. Method limitation: Although this method enables quantification of QS levels, it is limited to concentrations within a certain linear range.

### Outer membrane vesicle filtration

DMS3^vir-*aqs1* l^ysate from a PA14 Δ*pf5* or a PA14 Δpf5 Δ*lasI* Δ*rhlI* Δ*pqsA* propagation host was filtered on Amicon 10 kDa molecular weight cutoff centrifugal filters (Amicon®, UFC901024) and centrifuged at 1893 x g for 30 min, before storage at 4 °C in Falcon tubes. The retained membrane vesicles were washed twice in PBS and resuspended 1:1 of the original volume in PBS.

### Pyocyanin measurements

PA14 *pilA*::FRT and PA14 *pilA*::FRT Δ*pqsA* were grown ON from fresh colonies. The cultures were back diluted to an OD_600 o_f 0.1 in LB. 1.8 mL of the diluted culture was transferred to 14 mL Round-Bottom Polypropylene Tubes (352059, Falcon™). 20 µL, 60 µL, 100 µL, or 200 µL of DMS3^vir-*aqs1* l^ysate and/or LB was added to a final volume of 2 mL. All samples were incubated for 19 h with 300 RPM shaking at 37 °C. The tubes were photographed and the OD_600 w_as measured (Implen OD_600®_). The cultures were centrifuged at 16,000 x *g* for 3 min, and pyocyanin was quantified by measuring the OD_695 o_f the supernatants using a GENESYS™ 140/150 Vis/UV-Vis Spectrophotometer (Thermo Scientific™). The entire photograph of the cultures was adjusted uniformly for contrast in Microsoft Word.

### *G. mellonella* infection assay

*G. mellonella* wax moth larvae (raised antibiotic-free) were obtained from a pet shop and stored at 12 °C for up to 10 days prior to use. Larvae 2.5-3.0 cm in length and with no melanization were sterilized with 70% ethanol and briefly returned to 12 °C until further use (2 larvae per dish). Burn wounds were created by heating the head of a 2 mm diameter stainless-steel nail with a Bunsen burner until red-hot, allowing it to cool for 4 seconds, and gently applying it to the dorsal surface of each larva for 4 sec. Fresh wounds were infected by slowly pipetting 10 μL of an OD_600 =_ 1 standardized ON culture of *P. aeruginosa* PA14 WT onto the wound. Where indicated, larvae received 10⁹ PFU of either QS− or QS+ phage lysate onto the burn wound, administered either as the sole treatment or one h after bacterial infection. Larvae were incubated at 37 °C for up to 4 days and monitored for survival every 24 h.

### *pqsABCDE* reporter assay

*P. aeruginosa* PA14 *pqsABCDE*::*luxCDABE* was grown ON to an OD_600 ≈_ 5 and back diluted 10x. The diluted culture was treated with 10% of the total volume of LB or DMS3^vir-*aqs1* g^rown on either PA14 Δ*pf5* or from PA14 Δ*pf5* Δ*lasI* Δ*rhlI* Δ*pqsA.* The OD_600 a_nd luminescence were measured every 20 min for 24 h at 37 °C, with shaking. All measurements were made in technical triplicates using a Synergy H1 plate reader (BioTek Instruments Inc.) and software Gen5 v.3.05.11. The luminescence signals were divided by the corresponding OD_600 t_o obtain RLUs.

### Swarm assay

Petri dishes (10 cm diameter) containing 21 mL swarming media (M9 medium supplemented with 1 mM MgSO_4,_ 0.2% glucose, 0.5% Casamino Acids (Becton, Dickinson), 1 mg cholesterol, and 0.5% agar) were prepared. The solidified swarm plates were dried 40 min without lids in a LAF-bench. 5 µL PA14 ON culture was spotted at the center of the plate. For phage infection assays, 1 µL of DMS3^vir-*aqs1* f^rom either a PA14 Δ*pf5* or Δ*pf5* Δ*lasI* Δ*rhlI* Δ*pqsA* propagation host was mixed with 5 µL PA14 ON culture and spotted at satellite positions in a concentric circle around the center spot. For the phage lysate-only satellites, 1 µL of each lysate was mixed with 5 µL MilliQ and spotted. PQS (94398; Sigma-Aldrich) was diluted to the indicated concentrations in DMSO and 6 µl was spotted. Plates were incubated 16-20 h at 37 °C in a humid container.

### Statistical analysis

GraphPad Prism version 10.4.1 for Windows (GraphPad Software, Massachusetts USA) was used for analyses. An unpaired two-tailed t-test was used for comparisons between two samples, while a one-way ANOVA was performed when comparing three or more samples. For ANOVAs, post-hoc Tukey HSD was performed to correct for multiple testing. Samples with values below the linear ranges of the standard curves of the reporter strains (Fig. 1A+C+D) were not included in the statistical testing. Samples with only technical replicates were also not included in the statistical testing (Fig. 1B).

## Data availability

All data are included in the supplementary Excel sheet at the point of publication.

## Acknowledgements

We thank Albert Siryaporn, George O’Toole, Joseph Bondy-Denomy, Lars Hestbjerg Hansen, and Félix d’Hérelle Reference Center for Bacterial Viruses, the Université Laval for sharing bacterial strains and phages. The study was supported by the Independent Research Fund Denmark grant 1054-00099B to NMH-K, the Novo Nordisk Foundation grant NNF24OC0096270 to NMH-K and JJM, and a research grant from the Danish Advanced Research Academy, funded by the Novo Nordisk Foundation under grant number NNF25SA0100969 to AHE.

## Supplementary Files

**Supplemental Figure 1:**
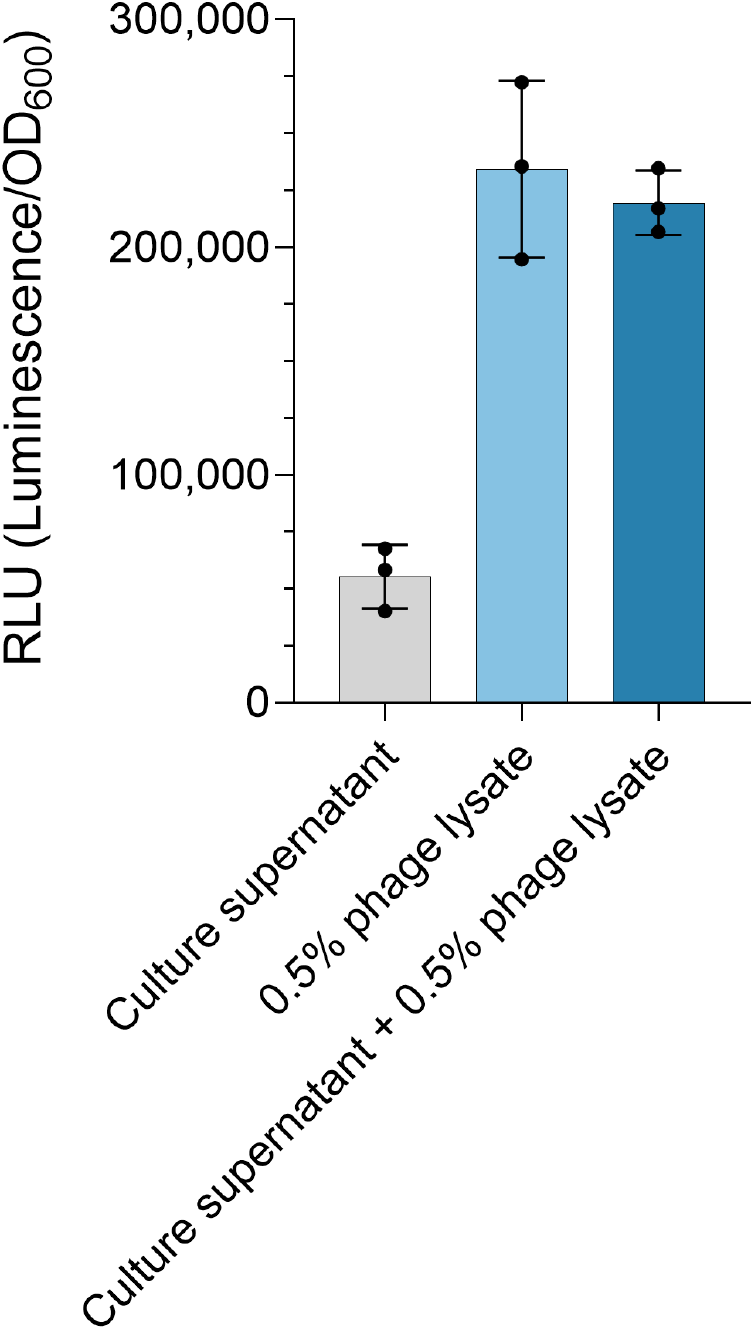
PQS is present in phage lysates. The relative PQS signal was measured in a DMS3^vir-*aqs1* p^hage lysate prepared on PA14, or in sterile filtered supernatant of a PA14 culture 2h after control (LB) treatment or infection with a multiplicity of infection (MOI) 10 DMS3^vir-*aqs1*.^ The PQS signals were determined using an *E. coli* PQS reporter strain. The phage lysate constituted 0.5% of the final volume in both the lysate-only control and the phage-infected culture. Relative light units (RLU) represent the luminescence of the PQS reporter strain normalized to OD_600._ Error bars indicate the standard deviation of three technical replicates.

**Supplemental Figure 2:**
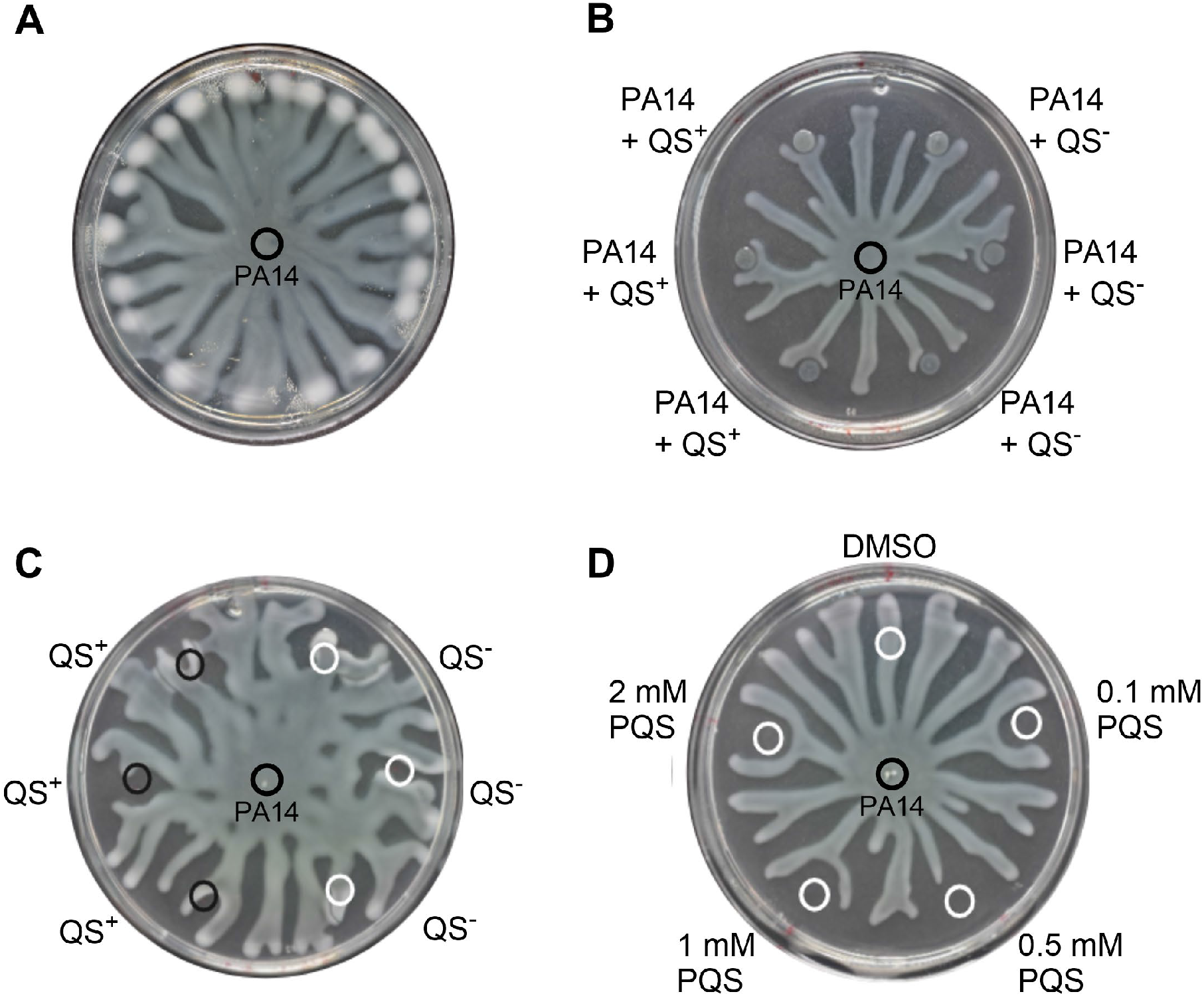
QS+ and QS-phage infection repels *P. aeruginosa* swarms. A. WT PA14 without phage B. Uninfected WT PA14 was spotted at the center, and PA14 infected with either QS- or QS+ phage lysates was spotted at satellite positions surrounding the center. C. Uninfected WT PA14 was spotted at the center, and either QS- or QS+ phage lysates were spotted at satellite positions surrounding the center. D. Uninfected WT PA14 was spotted at the center, with DMSO or increasing concentrations (mM) of PQS spotted at the satellite positions. Representative images are shown.

**Supplemental Table 1:**
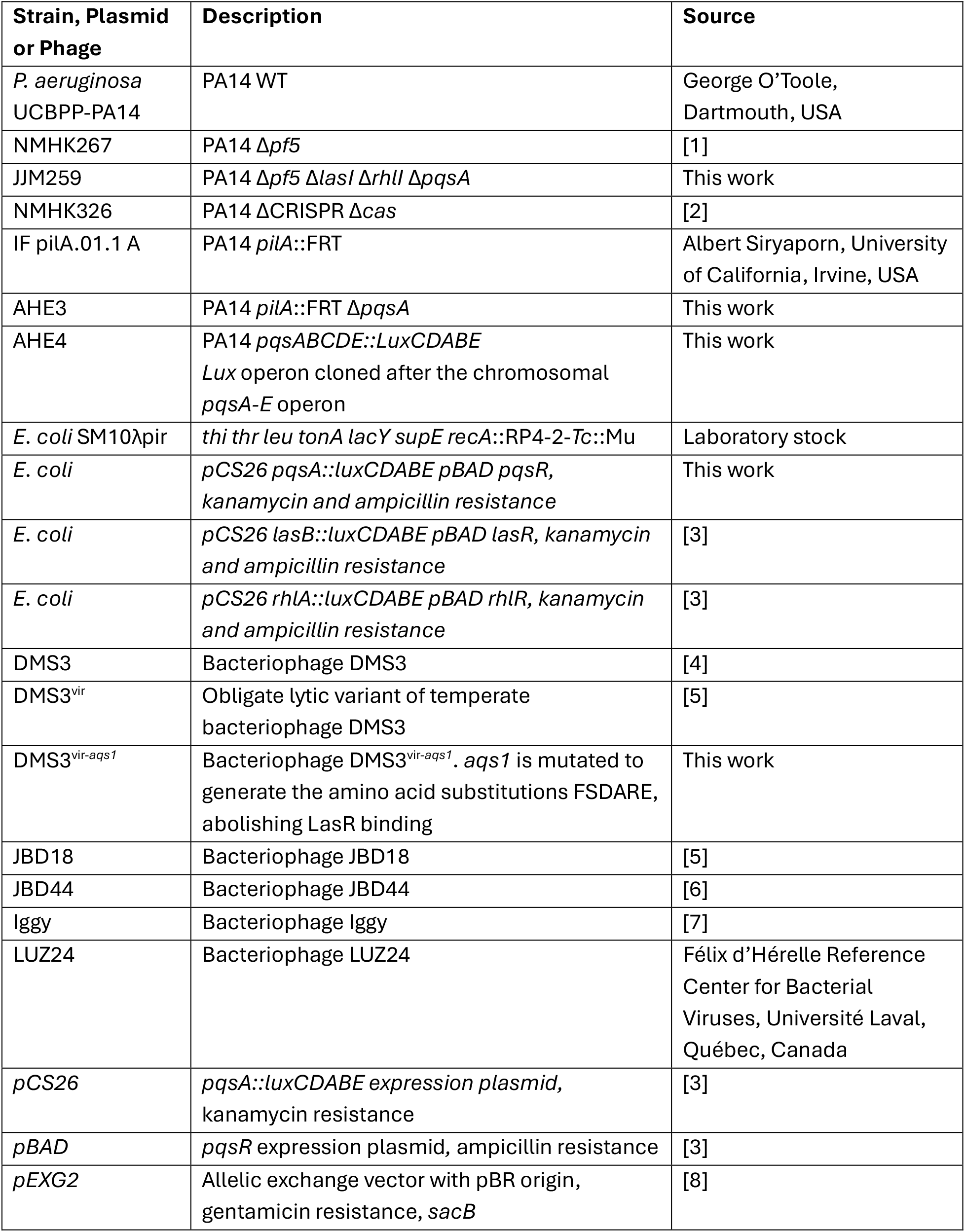
Bacterial strains, plasmids and bacteriophages.

| **Strain, Plasmid or Phage** | **Description** | **Source** |
|---|---|---|
| *P. aeruginosa* UCBPP-PA14 | PA14 WT | George O’Toole, Dartmouth, USA |
| NMHK267 | PA14 Δ*pf5* | [1] |
| JJM259 | PA14 Δ*pf5* Δ*lasI* Δ*rhlI* Δ*pqsA* | This work |
| NMHK326 | PA14 ΔCRISPR Δ*cas* | [2] |
| IF pilA.01.1 A | PA14 *pilA*::FRT | Albert Siryaporn, University of California, Irvine, USA |
| AHE3 | PA14 *pilA*::FRT Δ*pqsA* | This work |
| AHE4 | PA14 *pqsABCDE::LuxCDABE*<br>*Lux* operon cloned after the chromosomal *pqsA-E* operon | This work |
| *E. coli* SM10λpir | *thi thr leu tonA lacY supE recA*::RP4-2-*Tc*::Mu | Laboratory stock |
| *E. coli* | *pCS26 pqsA::luxCDABE pBAD pqsR, kanamycin and ampicillin resistance* | This work |
| *E. coli* | *pCS26 lasB::luxCDABE pBAD lasR, kanamycin and ampicillin resistance* | [3] |
| *E. coli* | *pCS26 rhlA::luxCDABE pBAD rhlR, kanamycin and ampicillin resistance* | [3] |
| DMS3 | Bacteriophage DMS3 | [4] |
| DMS3<sup>vir</sup> | Obligate lytic variant of temperate bacteriophage DMS3 | [5] |
| DMS3<sup>vir-*aqs1*</sup> | Bacteriophage DMS3<sup>vir-*aqs1*</sup>. *aqs1* is mutated to generate the amino acid substitutions FSDARE, abolishing LasR binding | This work |
| JBD18 | Bacteriophage JBD18 | [5] |
| JBD44 | Bacteriophage JBD44 | [6] |
| Iggy | Bacteriophage Iggy | [7] |
| LUZ24 | Bacteriophage LUZ24 | Félix d’Hérelle Reference Center for Bacterial Viruses, Université Laval, Québec, Canada |
| *pCS26* | *pqsA::luxCDABE expression plasmid,* kanamycin resistance | [3] |
| *pBAD* | *pqsR* expression plasmid, ampicillin resistance | [3] |
| *pEXG2* | Allelic exchange vector with pBR origin, gentamicin resistance, *sacB* | [8] |

**Supplemental Table 2:** Phage lysates used in this study.

| **Sample** | **Produced** | **Phage** | **Host** | **Preparation method** |
|---|---|---|---|---|
| **Fig. 1B - PQS measurements** |  |  |  |  |
| 1 week | 01/2025 | DMS3<sup>vir-*aqs1*</sup> | PA14 Δ*pf5* | Plate lysate |
| 3 months | 10/2024 | DMS3<sup>vir-*aqs1*</sup> | PA14 Δ*pf5* | Plate lysate |
| 1 year | 12/2023 | LUZ24 | PA14 WT | Plate lysate |
| 3 years A | 06/2021 | JBD18 | SMC5486*cas* | ON culture of lysogen |
| 3 years B | 06/2021 | JBD25 | SMC5487 ΔCRISPR Δ*cas* | ON culture of lysogen |
| 5 years | 08/ 2019 | Iggy | PA14 ΔCRISPR Δ*cas* | Plate lysate |
| 6 years A | 11/ 2018 | DMS3<sup>m</sup> | PA14 ΔCRISPR Δ*cas* | Plate lysate |
| 6 years B | 11/ 2018 | DMS3<sup>m</sup> | PA14 ΔCRISPR Δ*cas* | Plate lysate |
| 6 years C | 11/ 2018 | DMS3<sup>vir</sup> | PA14 ΔCRISPR Δ*cas* | Plate lysate |
| 6 years D | 11/ 2018 | DMS3<sup>mvir</sup> | PA14 ΔCRISPR Δ*cas* | Plate lysate |
| **Fig. 1C + 1D - AHL measurements** |  |  |  |  |
| **Sample** | **Produced** | **Phage** | **Host** | **Preparation method** |
| 1-4 days (QS+) | 10/2025 | DMS3<sup>vir-*aqs1*</sup> | PA14 Δ*pf5* | Plate lysate |
| 1-4 days (QS-) | 10/2025 | DMS3<sup>vir-*aqs1*</sup> | PA14 Δ*pf5* Δ*las*I Δ*rhlI* Δ*pqsA* | Plate lysate |

## References

Abdalla, M. Y., Hoke, T., Seravalli, J., Switzer, B. L., Bavitz, M., Fliege, J. D., Murphy, P. J., & Britigan, B. E. (2017). Pseudomonas quinolone signal induces oxidative stress and inhibits heme oxygenase-1 expression in lung epithelial cells. Infection and Immunity, 85(9). 10.1128/IAI.00176-17

Augustyniak, D., Olszak, T., & Drulis-Kawa, Z. (2022). Outer Membrane Vesicles (OMVs) of Pseudomonas aeruginosa Provide Passive Resistance but Not Sensitization to LPS-Specific Phages. Viruses, 14(1), 121. 10.3390/V14010121/S1

Bala, A., Gupta, R. K., Chhibber, S., & Harjai, K. (2013). Detection and quantification of quinolone signalling molecule: A third quorum sensing molecule of Pseudomonas aeruginosa by high performance-thin layer chromatography. Journal of Chromatography B: Analytical Technologies in the Biomedical and Life Sciences, 930, 30–35. 10.1016/j.jchromb.2013.04.027

Bala, A., Kumar, L., Chhibber, S., & Harjai, K. (2015). Augmentation of virulence related traits of pqs mutants by Pseudomonas quinolone signal through membrane vesicles. Journal of Basic Microbiology, 55(5), 566–578. 10.1002/JOBM.201400377

Blasdel, B. G., Ceyssens, P.-J., Chevallereau, A., Debarbieux, L., & Lavigne, R. (2018). Comparative transcriptomics reveals a conserved Bacterial Adaptive Phage Response (BAPR) to viral predation. BioRxiv, 248849. 10.1101/248849

Brameyer, S., Plener, L., Müller, A., Klingl, A., Wanner, G., & Jung, K. (2018). Outer membrane vesicles facilitate trafficking of the hydrophobic signaling molecule CAI-1 between Vibrio harveyi cells. Journal of Bacteriology, 200(15). 10.1128/JB.00740-17,

Bru, J.-L., Rawson, B., Trinh, C., Whiteson, K., Molin Høyland-Kroghsbo, N., & Siryaporn, A. (2019). PQS Produced by the Pseudomonas aeruginosa Stress Response Repels Swarms Away from Bacteriophage and Antibiotics. Journal of Bacteriology, (23). 10.1128/JB/00383-19

Budzik, J. M., Rosche, W. A., Rietsch, A., & O’Toole, G. A. (2004). Isolation and characterization of a generalized transducing phage for Pseudomonas aeruginosa strains PAO1 and PA14. Journal of Bacteriology, 186(10), 3270–3273. 10.1128/JB.186.10.3270-3273.2004

Byers, J. T., Lucas, C., Salmond, G. P. C., & Welch, M. (2002). Nonenzymatic turnover of an Erwinia carotovora quorum-sensing signaling molecule. Journal of Bacteriology, 184(4), 1163–1171. 10.1128/JB.184.4.1163-1171.2002

Déziel, E., Lépine, F., Milot, S., He, J., Mindrinos, M. N., Tompkins, R. G., & Rahme, L. G. (2004). Analysis of Pseudomonas aeruginosa 4-hydroxy-2-alkylquinolines (HAQs) reveals a role for 4-hydroxy-2-heptylquinoline in cell-to-cell communication. Proceedings of the National Academy of Sciences of the United States of America, 101(5), 1339–1344. 10.1073/PNAS.0307694100

Diggle, S. P., Winzer, K., Chhabra, S. R., Worrall, K. E., Cámara, M., & Williams, P. (2003). The Pseudomonas aeruginosa quinolone signal molecule overcomes the cell density-dependency of the quorum sensing hierarchy, regulates rhl-dependent genes at the onset of stationary phase and can be produced in the absence of LasR. Molecular Microbiology, 50(1), 29–43. 10.1046/J.1365-2958.2003.03672.X

Dulcey, C. E., Dekimpe, V., Fauvelle, D. A., Milot, S., Groleau, M. C., Doucet, N., Rahme, L. G., Lépine, F., & Déziel, E. (2013). The end of an old hypothesis: The pseudomonas signaling molecules 4-hydroxy-2-alkylquinolines derive from fatty acids, not 3-ketofatty acids. Chemistry and Biology, 20(12), 1481–1491. 10.1016/j.chembiol.2013.09.021

Erez, Z., Steinberger-Levy, I., Shamir, M., Doron, S., Stokar-Avihail, A., Peleg, Y., Melamed, S., Leavitt, A., Savidor, A., Albeck, S., Amitai, G., & Sorek, R. (2017). Communication between viruses guides lysis–lysogeny decisions. Nature 2017 541:7638, 541(7638), 488–493. 10.1038/nature21049

Florez, C., Raab, J. E., Cooke, A. C., & Schertzer, J. W. (2017). Membrane distribution of the Pseudomonas quinolone signal modulates outer membrane vesicle production in Pseudomonas aeruginosa. MBio, 8(4). 10.1128/mBio.01034-17

Gallagher, L. A., McKnight, S. L., Kuznetsova, M. S., Pesci, E. C., & Manoil, C. (2002). Functions required for extracellular quinolone signaling by Pseudomonas aeruginosa. Journal of Bacteriology, 184(23), 6472–6480. 10.1128/JB.184.23.6472-6480.2002/ASSET/ABE079E9-8B2D-453B-B626-3BA820288579/ASSETS/GRAPHIC/JB2320613004.JPEG

Glansdorp, F. G., Thomas, G. L., Lee, J. K., Dutton, J. M., Salmond, G. P. C., Welch, M., & Spring, D. R. (2004). Synthesis and stability of small molecule probes for Pseudomonas aeruginosa quorum sensing modulation. Organic & Biomolecular Chemistry, 2(22), 3329–3336. 10.1039/b412802h

Høyland-Kroghsbo, N. M., & Bassler, B. L. (2022). Phage Infection Restores PQS Signaling and Enhances Growth of a Pseudomonas aeruginosa lasI Quorum-Sensing Mutant. Journal of Bacteriology. 10.1128/jb.00557-21

Høyland-Kroghsbo, N. M., Mærkedahl, R. B., & Svenningsen, S. Lo. (2013). A quorum-sensing-induced bacteriophage defense mechanism. MBio, 4(1). 10.1128/MBIO.00362-12

Høyland-Kroghsbo, N. M., Paczkowski, J., Mukherjee, S., Broniewski, J., Westra, E., Bondy-Denomy, J., & Bassler, B. L. (2017). Quorum sensing controls the pseudomonas aeruginosa CRISPR-Cas adaptive immune system. Proceedings of the National Academy of Sciences of the United States of America, 114(1), 131–135. 10.1073/PNAS.1617415113

Latifi, A., Winson, M. K., Foglino, M., Bycroft, B. W., Stewart, G. S. A. B., Lazdunski, A., & Williams, P. (1995). Multiple homologues of LuxR and LuxI control expression of virulence determinants and secondary metabolites through quorum sensing in Pseudomonas aeruginosa PAO1. Molecular Microbiology, 17(2), 333–343. 10.1111/j.1365-2958.1995.mmi_17020333.x

Manning, A. J., & Kuehn, M. J. (2011). Contribution of bacterial outer membrane vesicles to innate bacterial defense. BMC Microbiology, 11(1), 1–15. 10.1186/1471-2180-11-258

Mashburn, L. M., & Whiteley, M. (2005). Membrane vesicles traffic signals and facilitate group activities in a prokaryote. Nature 2005 437:7057, 437(7057), 422–425. 10.1038/nature03925

Mashburn-Warren, L., Howe, J., Garidel, P., Richter, W., Steiniger, F., Roessle, M., Brandenburg, K., & Whiteley, M. (2008). Interaction of quorum signals with outer membrane lipids: Insights into prokaryotic membrane vesicle formation. Molecular Microbiology, 69(2), 491–502. 10.1111/j.1365-2958.2008.06302.x

Maslova, E., Shi, Y., Sjöberg, F., Azevedo, H. S., Wareham, D. W., & McCarthy, R. R. (2020). An Invertebrate Burn Wound Model That Recapitulates the Hallmarks of Burn Trauma and Infection Seen in Mammalian Models. Frontiers in Microbiology, 11, 527842. 10.3389/fmicb.2020.00998

Mauritzen, J. J., Koonce, kira C., Lassen, S., & Hoyland-Kroghsbo, N. M. (2026). Anti-Quorum Sensing Phages Disarm Pseudomonas aeruginosa. BioRxiv, 2026.02.16.706229. 10.64898/2026.02.16.706229

Morinaga, K., Nagakubo, T., Nomura, N., & Toyofuku, M. (2020). Involvement of membrane vesicles in long-chain-AHL delivery in Paracoccus species. Environmental Microbiology Reports, 12(3), 355–360. 10.1111/1758-2229.12843,

Paczesny, J., & Bielec, K. (2020). Application of Bacteriophages in Nanotechnology. Nanomaterials, 10(10), 1944. 10.3390/NANO10101944

Paczkowski, J. E., Mukherjee, S., McCready, A. R., Cong, J. P., Aquino, C. J., Kim, H., Henke, B. R., Smith, C. D., & Bassler, B. L. (2017). Flavonoids Suppress Pseudomonas aeruginosa Virulence through Allosteric Inhibition of Quorum-sensing Receptors. Journal of Biological Chemistry, 292(10), 4064–4076. 10.1074/JBC.M116.770552

Papenfort, K., & Bassler, B. L. (2016). Quorum-Sensing Signal-Response Systems in Gram-Negative Bacteria. Nature Reviews. Microbiology, 14(9), 576. 10.1038/NRMICRO.2016.89

Patterson, A. G., Jackson, S. A., Taylor, C., Evans, G. B., Salmond, G. P. C., Przybilski, R., Staals, R. H. J., & Fineran, P. C. (2016). Quorum Sensing Controls Adaptive Immunity through the Regulation of Multiple CRISPR-Cas Systems. Molecular Cell, 64(6), 1102. 10.1016/J.MOLCEL.2016.11.012

Pearson, J. P., Passador, L., Iglewski, B. H., & Greenberg, E. P. (1995). A second N-acylhomoserine lactone signal produced by Pseudomonas aeruginosa. Proceedings of the National Academy of Sciences, 92(5), 1490–1494. 10.1073/PNAS.92.5.1490

Pesci, E. C., Pearson, J. P., Seed, P. C., & Iglewski, B. H. (1997). Regulation of las and rhl quorum sensing in Pseudomonas aeruginosa. Journal of Bacteriology, 179(10), 3127. 10.1128/JB.179.10.3127-3132.1997

Reyes-Robles, T., Dillard, R. S., Cairns, L. S., Silva-Valenzuela, C. A., Housman, M., Ali, A., Wright, E. R., & Camilli, A. (2018). Vibrio cholerae Outer Membrane Vesicles Inhibit Bacteriophage Infection. Journal of Bacteriology, 200(15), e00792–17. 10.1128/JB.00792-17

Rietsch, A., Vallet-Gely, I., Dove, S. L., & Mekalanos, J. J. (2005). ExsE, a secreted regulator of type III secretion genes in Pseudomonas aeruginosa. Proceedings of the National Academy of Sciences of the United States of America, 102(22), 8006–8011. 10.1073/PNAS.0503005102

Rueter, C., & Bielaszewska, M. (2020). Secretion and Delivery of Intestinal Pathogenic Escherichia coli Virulence Factors via Outer Membrane Vesicles. Frontiers in Cellular and Infection Microbiology, 10, 519869. 10.3389/FCIMB.2020.00091/EPUB

Schertzer, J. W., Brown, S. A., & Whiteley, M. (2010). Oxygen levels rapidly modulate Pseudomonas aeruginosa social behaviours via substrate limitation of PqsH. Molecular Microbiology, 77(6), 1527–1538. 10.1111/j.1365-2958.2010.07303.x

Schwartzkopf, C. M., Taylor, V. L., Groleau, M. C., Faith, D. R., Schmidt, A. K., Lamma, T. L., Brooks, D. M., Déziel, E., Maxwell, K. L., & Secor, P. R. (2023). Inhibition of PQS signaling by the Pf bacteriophage protein PfsE enhances viral replication in Pseudomonas aeruginosa. Molecular Microbiology, 121(1), 116. 10.1111/MMI.15202

Seed, P. C., Passador, L., & Iglewski, B. H. (1995). Activation of the Pseudomonas aeruginosa lasI gene by LasR and the Pseudomonas autoinducer PAI: an autoinduction regulatory hierarchy. Journal of Bacteriology, 177(3), 654. 10.1128/jb.177.3.654-659.1995

Shah, M., Taylor, V. L., Bona, D., Tsao, Y., Stanley, S. Y., Pimentel-Elardo, S. M., McCallum, M., Bondy-Denomy, J., Howell, P. L., Nodwell, J. R., Davidson, A. R., Moraes, T. F., & Maxwell, K. L. (2021). A phage-encoded anti-activator inhibits quorum sensing in Pseudomonas aeruginosa. Molecular Cell, 81(3), 571–583.e6. 10.1016/j.molcel.2020.12.011

Silpe, J. E., & Bassler, B. L. (2018). A Host-Produced Quorum-Sensing Autoinducer Controls a Phage Lysis-Lysogeny Decision. Cell, 176(1–2), 268. 10.1016/J.CELL.2018.10.059

Tan, D., Svenningsen, S. Lo, & Middelboe, M. (2015). Quorum Sensing Determines the Choice of Antiphage Defense Strategy in Vibrio anguillarum. MBio, 6(3). 10.1128/MBIO.00627-15

Turner, P. E., Azeredo, J., Buurman, E. T., Green, S., Haaber, J. K., Haggstrom, D., Kameda de Figueiredo Carvalho, K., Kirchhelle, C., Gonzalez Moreno, M., Pirnay, J.-P., & Portillo, M. A. (2024). Addressing the Research and Development Gaps in Modern Phage Therapy. PHAGE: Therapy, Applications, and Research, 5(1), 30–39. 10.1089/phage.2023.0045

Wade, D. S., Calfee, M. W., Rocha, E. R., Ling, E. A., Engstrom, E., Coleman, J. P., & Pesci, E. C. (2005). Regulation of *Pseudomonas* Quinolone Signal Synthesis in *Pseudomonas aeruginosa*. Journal of Bacteriology, 187(13), 4372–4380. 10.1128/JB.187.13.4372-4380.2005

Xiao, G., Déziel, E., He, J., Lépine, F., Lesic, B., Castonguay, M. H., Milot, S., Tampakaki, A. P., Stachel, S. E., & Rahme, L. G. (2006). MvfR, a key Pseudomonas aeruginosa pathogenicity LTTR-class regulatory protein, has dual ligands. Molecular Microbiology, 62(6), 1689–1699. 10.1111/j.1365-2958.2006.05462.x

Xiao, G., He, J., & Rahme, L. G. (2006). Mutation analysis of the Pseudomonas aeruginosa mvfR and pqsABCDE gene promoters demonstrates complex quorum-sensing circuitry. Microbiology, 152(6), 1679–1686. 10.1099/mic.0.28605-0

Yates, E. A., Philipp, B., Buckley, C., Atkinson, S., Chhabra, S. R., Sockett, R. E., Goldner, M., Dessaux, Y., Cámara, M., Smith, H., & Williams, P. (2002). N-acylhomoserine lactones undergo lactonolysis in a pH-, temperature-, and acyl chain length-dependent manner during growth of Yersinia pseudotuberculosis and Pseudomonas aeruginosa. Infection and Immunity, 70(10), 5635–5646. 10.1128/IAI.70.10.5635-5646.2002

## References

[1] N. M. Høyland-Kroghsbo and B. L. Bassler, “Phage Infection Restores PQS Signaling and Enhances Growth of a Pseudomonas aeruginosa lasI Quorum-Sensing Mutant,” J. Bacteriol., 2022, doi: 10.1128/jb.00557-21.

[2] N. M. Høyland-Kroghsbo et al., “Quorum sensing controls the pseudomonas aeruginosa CRISPR-Cas adaptive immune system,” Proc. Natl. Acad. Sci. U. S. A., vol. 114, no. 1, pp. 131–135, Jan. 2017, doi: 10.1073/PNAS.1617415113.

[3] J. E. Paczkowski et al., “Flavonoids Suppress Pseudomonas aeruginosa Virulence through Allosteric Inhibition of Quorum-sensing Receptors,” Journal of Biological Chemistry, vol. 292, no. 10, pp. 4064–4076, Mar. 2017, doi: 10.1074/JBC.M116.770552.

[4] J. M. Budzik, W. A. Rosche, A. Rietsch, and G. A. O’Toole, “Isolation and characterization of a generalized transducing phage for Pseudomonas aeruginosa strains PAO1 and PA14,” J. Bacteriol., vol. 186, no. 10, pp. 3270–3273, May 2004, doi: 10.1128/JB.186.10.3270-3273.2004.

[5] K. C. Cady, J. Bondy-Denomy, G. E. Heussler, A. R. Davidson, and G. A. O’Toole, “The CRISPR/Cas adaptive immune system of Pseudomonas aeruginosa mediates resistance to naturally occurring and engineered phages,” J. Bacteriol., vol. 194, no. 21, pp. 5728–5738, Nov. 2012, doi: 10.1128/JB.01184-12.

[6] A. Phee, J. Bondy-Denomy, A. Kishen, B. Basrani, A. Azarpazhooh, and K. Maxwell, “Efficacy of bacteriophage treatment on Pseudomonas aeruginosa biofilms,” J. Endod., vol. 39, no. 3, pp. 364–369, Mar. 2013, doi: 10.1016/J.JOEN.2012.10.023.

[7] N. S. Olsen et al., “A novel Queuovirinae lineage of Pseudomonas aeruginosa phages encode dPreQ0 DNA modifications with a single GA motif that provide restriction and CRISPR Cas9 protection in vitro,” Nucleic Acids Res., vol. 51, no. 16, pp. 8663–8676, Sep. 2023, doi: 10.1093/NAR/GKAD622.

[8] A. Rietsch, I. Vallet-Gely, S. L. Dove, and J. J. Mekalanos, “ExsE, a secreted regulator of type III secretion genes in Pseudomonas aeruginosa,” Proc. Natl. Acad. Sci. U. S. A., vol. 102, no. 22, pp. 8006–8011, May 2005, doi: 10.1073/PNAS.0503005102.

